# Early neuronal processes interact with glia to establish a scaffold for orderly innervation of the cochlea

**DOI:** 10.1101/754416

**Authors:** N. R. Druckenbrod, E. B. Hale, O. O. Olukoya, W. E. Shatzer, L.V. Goodrich

## Abstract

Although the basic principles of axon guidance are well established, it remains unclear how axons navigate with high fidelity through the complex cellular terrains that are encountered *in vivo*. To learn more about the cellular strategies underlying axon guidance *in vivo*, we analyzed the developing cochlea, where spiral ganglion neurons extend processes through a heterogeneous cellular environment to form tonotopically ordered connections with hair cells. Here, we show that the earliest processes are closely associated with a population of glia that grow ahead of them. By analyzing single cell morphology and imaging the real time behavior of neuronal processes and glia in embryonic cochleae, we show that spiral ganglion neurons employ different mechanisms depending on their position in the ganglion. Additionally, the pattern of outgrowth varied locally, with evidence for both glia-guided growth and fasciculation along a neuronal scaffold. These findings suggest a tiered mechanism for reliable axon guidance.

## Introduction

Neuroscientists have long puzzled how neurons make the connections needed for proper circuit function, with early mechanical explanations by Weiss ^1^ (1941) eventually set aside in favor of Sperry’s chemoaffinity hypothesis ^2^. With the discovery of axon guidance molecules, the field coalesced around the idea that axons are guided towards their targets by a combination of attractive and repulsive cues that act at short or long range, with direction specified by target-derived gradients ^3^. However, mathematical modeling studies suggest that chemoattractive gradients are not solely responsible for the remarkable precision of guidance events *in vivo*, where axons grow through complex and changing environments ^4^. Although synergy among cues may improve fidelity, other cellular mechanisms likely contribute, such as fasciculation with pioneers and avoidance of repellant boundaries ^5^. For instance, in the spinal cord, commissural axons grow along a permissive substrate of Netrin-1 in the subpial region before turning towards an instructive gradient of Netrin-1 and other cues emanating from the floor plate ^6–9^, underscoring the idea that axons rely on different mechanisms as they move through distinct cellular landscapes.

The cochlea presents a distinct landscape for axon growth compared to the spinal cord, with neural processes organized into a spatial stereotyped pattern within a highly heterogeneous cellular environment. The cochlea is comprised of three fluid-filled ducts: scala vestibuli, scala media, and scala tympani (Fig. 1a). The auditory sensory epithelium, the organ of Corti, sits on the floor of scala media and vibrates in response to wavelengths of sound, thereby activating sensory hair cells. Information is transmitted to the central nervous system by the spiral ganglion neurons (SGNs), whose cell bodies sit just outside of the cochlea. SGNs are bipolar neurons that extend a peripheral process to innervate hair cells in the organ of Corti and a central process that innervates target neurons in the auditory brainstem. In the cochlea, hair cells and SGNs are arranged tonotopically, from high frequencies in the base to low frequencies in the apex (Fig. 1b). As such, tonotopy is represented in the spatial arrangement of the SGN peripheral processes, which are bundled together like spokes of a wheel. These radial bundles of SGN processes are separated from each other by the mesenchymal cells of the osseous spiral lamina (OSL) (Fig. 1c). Type I SGNs, which comprise ∼95% of the population ^10^, extend radial fibers that form synapses with the inner hair cells (IHCs). Type I SGN peripheral processes are ensheathed by neural crest-derived Schwann cells up until the habenula perforata, which is a series of holes through which the processes penetrate to reach the organ of Corti. Even the SGN cell bodies are myelinated, in this case by satellite glia ^11^. The remaining Type II SGNs extend unmyelinated processes that spiral among the outer hair cells (OHCs). Type I SGNs are primarily response for encoding sound information, while Type II SGNs have been proposed to play a role in sensing damage ^12^.

**Figure 1:**
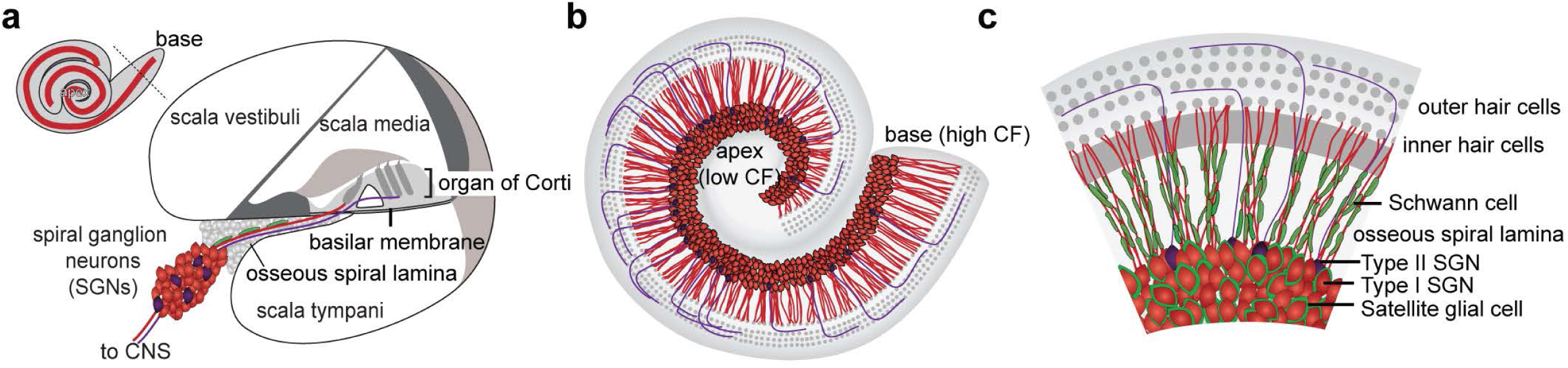
Overview of spiral ganglion neuron organization. (**a**) A schematic view of a cross section through the base of the cochlea. The cochlear duct is divided into three fluid-filled chambers (scala vestibuli, scala media, and scala tympani). The organ of Corti is overlaid by the tectorial membrane and sits on the basilar membrane, which divides scala media from scala tympani. Spiral ganglion neurons (SGNs, red and purple) sit outside the cochlear duct and extend peripheral processes through the osseous spiral lamina (gray) to reach the organ of Corti. Their central axons project through the eighth nerve into the central nervous system (CNS). (**b,c**) Low (b) and high-power (c) top-down views of the cochlear duct. SGNs extend peripheral processes through the osseous spiral lamina, and towards the organ of Corti (gray). Radial bundles of processes are spatially organized along the length of the cochlea, with low characteristic frequencies (CF) detected in the apex and high CFs in the base. Type I SGNs (red) project to the inner hair cells, and Type II SGNs (purple) project to outer hair cells, where they turn and spiral towards the base. Type I SGN cell bodies and peripheral processes are myelinated by satellite glial cells and Schwann cells respectively.

Unlike other regions of the nervous system, target-derived cues seem to play a minimal role in gross cochlear wiring ^13, 14^. For instance, in mice with no differentiated hair cells, such as *Atoh1* mutants, SGN peripheral processes still form radial bundles and reach the organ of Corti ^15, 16^. Additionally, although the classic chemoattractrant Netrin-1 can elicit outgrowth *in vitro* ^17^, it is not expressed in the organ of Corti and is not required for guidance towards the cochlea *in vivo* ^18^. On the other hand, there is a prominent role for repulsion ^19–22^. In the OSL, Ephrin-Eph signaling between the neurons and the mesenchyme ensures tight fasciculation of SGN processes in radial bundles ^21^, whereas in the organ of Corti, Semaphorin 3A and Ephrin A5 keep the processes from entering the OHC region ^20, 22^. Thus, once the processes start growing, they may be kept on course by repulsive cues encountered along the way. How peripheral axons begin this journey, however, remains unknown.

Several observations suggest that glia might be involved in the earliest stages of cochlear wiring. Early histological studies showed that SGN peripheral processes pass through a “glial-funnel” ^23^. Likewise, three-dimensional reconstructions revealed interdigitation of glia with the earliest growing neurites in the otic vesicle ^24^. There are also hints that glia are necessary for normal cochlear innervation. For example, in mice with impaired invasion of the cochlea by neural crest cell-derived glia, radial bundles do not form normally ^25, 26^. However, in these animals, the SGNs are also mispositioned, making it hard to know whether the abnormal radial outgrowth is secondary to an earlier migration phenotype.

Here, we sought to define the cellular mechanisms underlying the earliest stages of SGN peripheral process growth, including the potential role for glia. Using a combination of time-lapse imaging and three-dimensional reconstructions of individual SGN processes in the context of the intact developing cochlea, we provide evidence that glia guide pioneer axons that establish a scaffold for the radial bundle organization of the mature cochlea. These findings highlight the multi-stepped nature of axon guidance *in vivo* and have important implications for efforts to re-wire the damaged cochlea.

## Results

### Early SGN peripheral processes extend along glia that precede them

Cochlear wiring occurs in a complex and dynamic environment, with Type I and Type II SGN processes growing towards the organ of Corti even as the cochlea itself lengthens and coils ^27^. SGNs originate in the otic vesicle between E9 and E12 in mouse and delaminate into the surrounding mesenchyme ^28, 29^. Differentiating SGNs extend peripheral processes back towards the organ of Corti, reaching nascent hair cells around E15 ^30^ and forming synapses by birth ^31^. There is a topographic gradient of development, with SGNs in the base maturing before those in the apex. Neural-crest derived Schwann cells and satellite glia are present from the earliest stages of neurite outgrowth ^24^. Thus, long before reaching target hair cells in the organ of Corti, the SGN peripheral processes interact with other neurons and their processes, as well as with glia and mesenchyme.

To gain insights into the cellular mechanisms that might elicit outgrowth of SGN peripheral processes towards the cochlear duct, we characterized the wavefront of peripheral process growth relative to glia, using anti-β-III tubulin or anti-Neurofilament antibodies to label the SGN processes and a *PLP-GFP* transgenic reporter ^32^ to label the glia. We focused on E14-E14.5, since this is when most processes cross the border of the ganglion ^21^, with radial bundles apparent around E15-E15.5 ^30^. Although SGN processes eventually extend into the developing OSL, we found that their earliest interactions are instead with the neural-crest derived glia, which differentiate into satellite glia and Schwann cells. In addition to the expected intermingling of glia and SGN cell bodies within the ganglion, the SGN peripheral processes lie on top of a distinct population of glia that protrudes from the ganglion and forms a bridge to the cochlear duct (Fig. 2a-c). This population stands out both for its morphology, with large flat cells, and for its more intense PLP-GFP signal.

**Figure 2:**
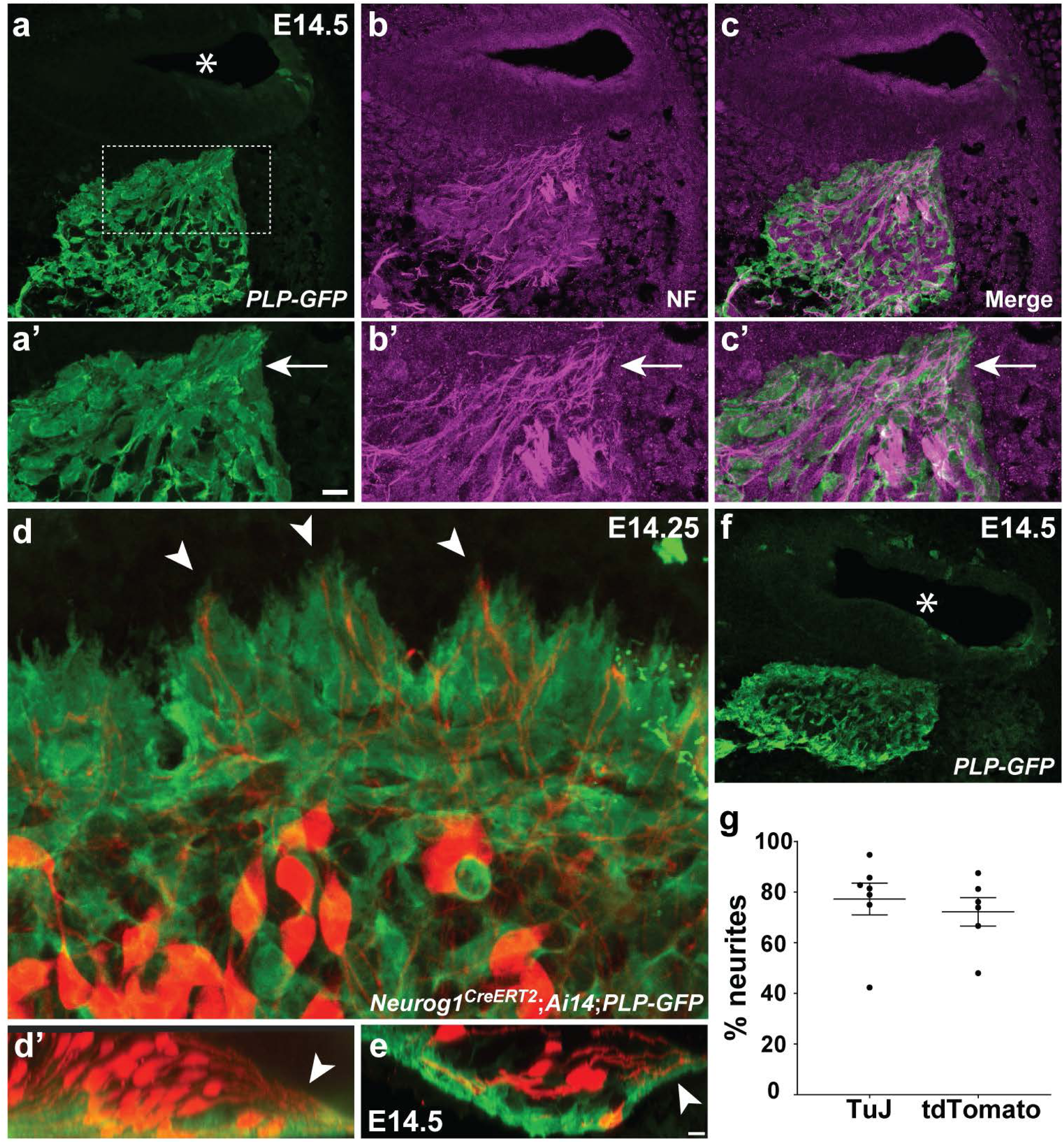
Distal SGN peripheral processes are closely associated with glia. (**a-c**) A cross-section through a head from an E14.5 PLP-GFP animal stained for GFP (green, a) and neurofilament (NF) (magenta, b), with a merged image shown in c. The lumen of the cochlear duct is indicated by an asterisk. Higher power views off the boxed region in a are shown in a’-c’. A swath of PLP-GFP+ glia forms a path to the cochlea for NF+ neurites (arrows, a’-c’). Behind this swath, the glia are intermingled with SGN cell bodies, as evidenced by the gaps in PLP-GFP signal in a and a’. (**d**) A high power top-down view of the wavefront of peripheral process growth in a wholemount E14.25 cochlea from a *Neurog1^CreERT2^;Ai14;PLP-GFP* animal that was stained for tdTomato (red) and GFP (green). The organ of Corti is up and the ganglion is down (as in Fig. 1c). Arrowheads indicate three examples of tdTomato-labeled peripheral processes that are preceded by glia. A side view of the same cochlea (d’) highlights the close relationship between the distalmost SGN peripheral processes (arrowhead) and a carpet of PLP-GFP^+^ glia, also evident in a single confocal slice from a stack through an E14.5 *Neurog1^CreERT2^;Ai14;PLP-GFP* wholemount cochlea (**e**). (**f**) A distinct population of PLP-GFP^+^ glia surround the ganglion, as viewed in a cross-section through an E14.5 head from a *PLP-GFP* animal that was stained for GFP (green). The lumen of the cochlear duct is indicated with an asterisk. (**g**) SGN peripheral processes were scored for whether they were ahead of or behind a glial cell in E14-E14.5 *PLP-GFP* cochlea stained for TuJ and GFP (N=7 cochleae; 164 neurites total) or E14-E14.5 *Neurog1^CreERT2^;Ai14;PLP-GFP* cochlea stained for tdTomato and GFP (N=6 cochleae; 125 neurites total). The percent of neurites that are preceded by a glial cell is plotted for each of the cochleae. Regardless of how SGN neurites were labeled, the vast majority of them grow behind a glial cell. There is no statistical difference between the percentages obtained using each method (*P*=0.5692, unpaired, two-tailed t-test).

To further define the relationship between the extending peripheral processes and this population of glia, we imaged SGN processes and glia in E14.25-E14.5 cochleae from *Neurog1^CreE1RT2^;Ai14*;*PLP-GFP* animals, in which a random subset of SGNs express tdTomato. At this early stage of development, no peripheral processes have grown more than 50 µm past the border of the ganglion. Wholemount preparations confirmed that the earliest extending SGN peripheral processes sit on top of a carpet of intensely PLP-GFP-positive glia (Fig. 2d, d’, e). Indeed, a population of strongly stained glia seem to form a shell around the entire ganglion, though it is possible that the intensity of the signal is simply higher because they are not interspersed with neuronal cell bodies here (Fig. 2f). Quantification showed that the glia are consistently ahead of the wavefront of peripheral process outgrowth: 72.25 % ± 5.63 (s.e.m.) of the most distal tdTomato-labeled SGN peripheral in the E14 cochlea were preceded by a PLP-GFP positive glial cell (n=125 neurites from 6 cochleae) (Fig. 2g). A similar relationship was observed when β-III tubulin+ peripheral processes were analyzed (77.28 % ± 6.27 s.e.m.) (n=164 neurites from 7 cochleae). These observations extend previous reports that neural crest-derived glia are closely affiliated with SGN and VGN processes during early stages of inner ear innervation ^24^, as well as the classic observation of a glial “funnel” through which early SGN processes appear to be steered towards the otic vesicle ^23^. Thus, glia are poised to influence the initial outgrowth of SGN peripheral processes.

### Spiral ganglion neurons exhibit heterogeneous morphologies and neurite outgrowth behavior in the early cochlea

Although the earliest SGN peripheral processes seem closely aligned with the glial carpet, other processes were positioned further away, raising the question of whether all SGNs follow the same path. To learn more about the range of trajectories taken by different SGNs, we made three-dimensional models of 143 SGNs from the mid-base of the cochlea in E14-E14.5 *Neurog1^CreE1RT2^;Ai14* animals (n=7) (Fig. 3a,b; Supp. Fig. 1). Consistent with impression that SGNs may grow differently depending on their local environment, we observed a range of morphologies that correlated with the position of each neuron’s cell body within the ganglion. SGNs whose cell bodies were close to the border extended twisted, short neurites (orange, Fig. 3b’, Video 1). By contrast, those whose cell bodies were in the rear of the ganglion exhibited long and directed neurites (purple, Fig. 3b’). SGNs in the middle of the ganglion showed intermediate morphologies (green, Fig. 3b’). Additionally, rear SGNs extended peripheral processes along a flat trajectory and were at the bottom of the pile of processes, closest to where the glial carpet is positioned (Fig. 3b”). By contrast, processes from the SGNs at the border were positioned at a steeper slope, diving down to meet the processes from the middle and rear SGNs (Fig. 3b”). These qualitative observations were confirmed by quantitative analysis of SGNs whose cell bodies sat near the border (“border cells”), in the middle region (“middle cells”), or in the rear (“rear cells”) (Fig. 3c-e and Supp. Fig. 2), using the peripheral circumferential border of the ganglion as a reference (Fig. 3a). The border cells showed consistently less directed, shorter processes that were at a steeper slope than those from the rear cells, with the middle cells falling in between. Thus, in the developing cochlea, SGN peripheral processes follow distinct paths depending on the position of their cell body in the ganglion: the rear SGN processes are closest to the glial carpet, with middle and border SGN processes layering on top.

**Figure 3:**
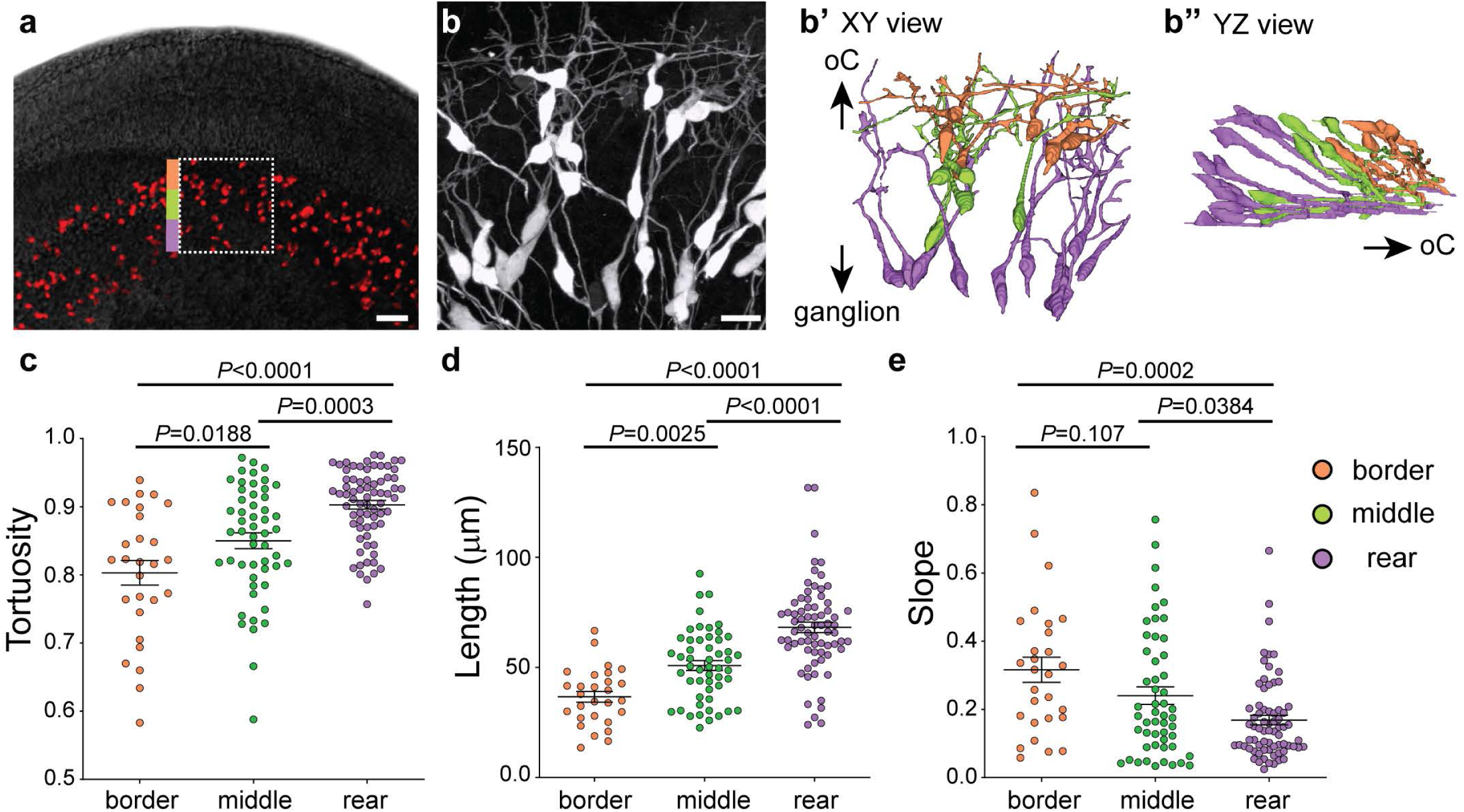
SGN morphology varies with cell body position. (**a**) A top-down view of a wholemount cochlea from an E14.5 *Neurog1^CreERT2^;Ai14* animal, where a random subset of SGNs express tdTomato (red). SGN morphology was analyzed at three positions in the ganglion: close to the border (orange), in the middle (green), or in the rear (purple). (**b**) A high power view of the boxed area in a, with reconstructions of individual SGNs shown in b’ (XY view) and b” (YZ view). SGNs are color coded according to the position of their cell body. (**c-e**) Quantification of SGN morphology. Border cells (orange dots) extend processes that are less directional (c), shorter (d), and have a greater slope (e) than rear cells. Middle cells exhibit intermediate morphologies. Significance was assessed using ANOVA with Tukey’s multiple comparison test; *P* values are indicated in each panel. Scale bars: 50 µm (a) and 25 µm (b). See Video 1.

Time lapse imaging in embryonic cochlear explants confirmed that SGN peripheral processes change their outgrowth behavior as they progress from the ganglion toward the developing organ of Corti. Since this entire journey take place from E14 to E15.5 *in vivo*, we imaged the overall wavefront of outgrowth at three separate positions along the trajectory: as the peripheral processes first exit the ganglion (at E14.25), as they are pushing through the OSL (around E15) and upon reaching the organ of Corti (at E15.5). At each stage, we analyzed outgrowth across a 50 µm region at the wavefront: immediately adjacent to the ganglion (R1), within the developing OSL (R2), and near the nascent organ of Corti (R3) (Fig 4a and Videos 2-4). The position of the tip of each process was plotted every 7-14 minutes (Fig. 4b,c), such that we could calculate the speed and direction of movement from the origin to the final position (n=3 cochleae) (Fig. 4d,e). We found that the speed and directionality of SGN process outgrowth varied along the trajectory. In R1, SGN peripheral processes (n=55) made slow progress (Fig 2d) and followed tortuous paths with many changes in direction (Fig 4e). By contrast, SGN process outgrowth was faster and highly directed within R2 (n=64), before slowing down and becoming less directed again in R3 (n=44). The behavior in R3 was similar to the exploratory behavior we observed previously in analysis of SGN peripheral processes close to the organ of Corti at E16.5^19^. Thus, even though the peripheral processes follow a relatively straight path towards the organ of Corti, they do not grow in the same way at each point along the trajectory.

**Figure 4:**
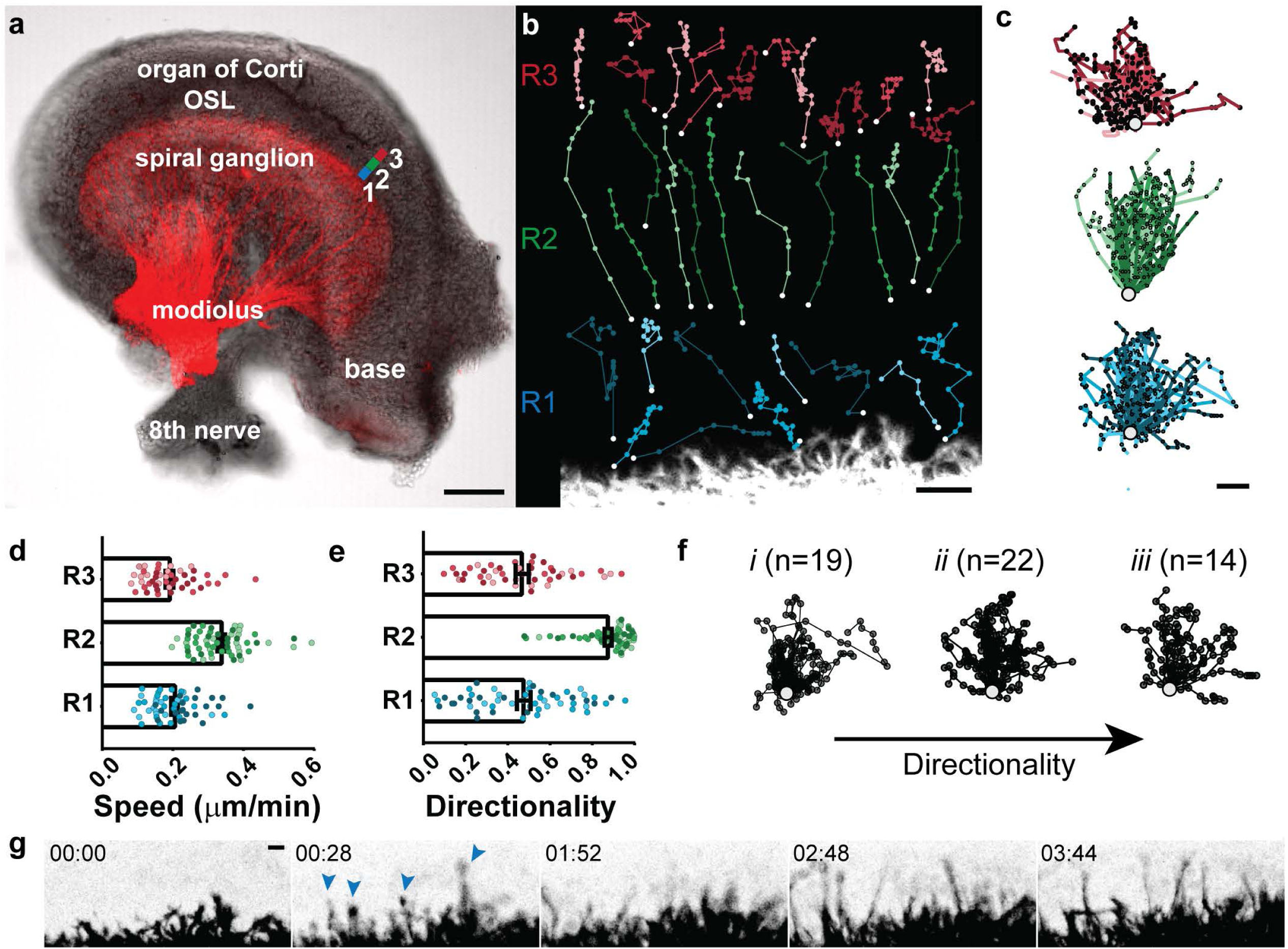
SGN peripheral processes exhibit a variety of outgrowth behaviors en route to the organ of Corti. (**a**) A wholemount E14 cochlea, with SGNs genetically labeled red using *Bhlhb5^cre^* and an *Ai14* tdTomato reporter. Process outgrowth was imaged in three different regions of E14-E15 cochleae. Region 1 (blue, R1) is immediately adjacent to the ganglion; Region 2 (green, R2) is in the developing osseous spiral lamina (OSL), and Region 3 (red, R3) is close to the organ of Corti. (**b**) Representative trajectories of processes imaged as they grew through each region. (**c**) All of the trajectories were collapsed onto a common origin, revealing overall differences in the pattern of growth in each region. N=3 cochleae for each region with n=55 R1 trajectories, 64 R2 trajectories, and 44 R3 trajectories. (**d,e**) Quantification of the speed (d) and directionality (e) of SGN peripheral process outgrowth in R1, R2, and R3. Growth is faster and more directed in R2 than in R1 or R3. Within R1, some SGN processes are as directed as those in R2, with directionality indices greater than 0.66. Significance was assessed using ANOVA with Tukey’s multiple comparison test. R1 vs R2 speed, *P*<0.0001; R1 vs R2 directionality, *P*<0.0001; R1 vs R3 directionality, *P*=0.9782; R1 vs R3 speed, *P*=0.6107; R2 vs R3 directionality, *P*<0.0001; and R2 vs R3 speed, *P*<0.0001. (**f**) R1 trajectories were grouped according to directionality, from low (0-0.33) (i) to medium (0.33-0.66) (ii) to high (0.66-1) (iii). The number of trajectories in group is indicated. (**g**) Frames from a video (Video 1) showing a subset of SGN peripheral processes (arrowheads) that grow ahead of the overall wavefront. Time is indicated in hh:mm. Scale bars: 100 µm (a), 10 µm (b, c, g). See Videos 2-4.

Closer examination of R1 revealed that even within a single location, there was heterogeneity in the behavior of the earliest extending SGN peripheral processes (Fig. 4f). While many processes showed slow, exploratory behavior consistent with the average behavior of the group, others grew in a fast, directed manner, occasionally darting ahead of the overall wavefront of neurite outgrowth (Fig. 4g and Video 2). Indeed, a subset of processes (14 neurites of 55 total, 25.5%) showed directionality similar to that in R2 (Fig. 4e and Fig. 4fiii). The remaining neurites followed more convoluted paths as they left the ganglion (Fig. 4fi,ii). Since SGN process morphology and trajectory varies with position (Fig. 3), these observations suggest that processes respond to local cues in their environment that affect how they exit the ganglion and grow through the mesenchyme.

### Processes from early born SGNs in the rear of the ganglion behave differently than follower neurites from SGNs at the border

Since analysis of morphologies revealed systematic differences among rear, border, and middle SGNs, we hypothesized that SGN peripheral processes may grow differently depending on where they are located in the ganglion and thus what kind of environment they first encounter. To characterize the behavior of individual axons extending from the rear or border of the ganglion, we performed time lapse imaging of E14-E14.5 cochleae from *Neurog1^CreERT^*^2^*;Ai14* animals. Since only some SGNs are labeled in this strain, we were able to define the cell body position for each extending peripheral process in R1. Consistent with the position-dependent variation in SGN morphology, border and rear SGNs showed strikingly different patterns of peripheral process outgrowth (Video 5). The peripheral processes from the rear SGNs followed fast, directed paths (blue arrows, Fig. 5a). By contrast, border SGNs instead showed exploratory, undirected outgrowth (orange dots, Fig. 5a). Quantification confirmed that trajectories from rear cells were significantly more directional than those from border cells (P<0.0001 by unpaired, two-tailed t-test; n=25 border cells and 24 rear cells from N=3 cochleae).

**Figure 5:**
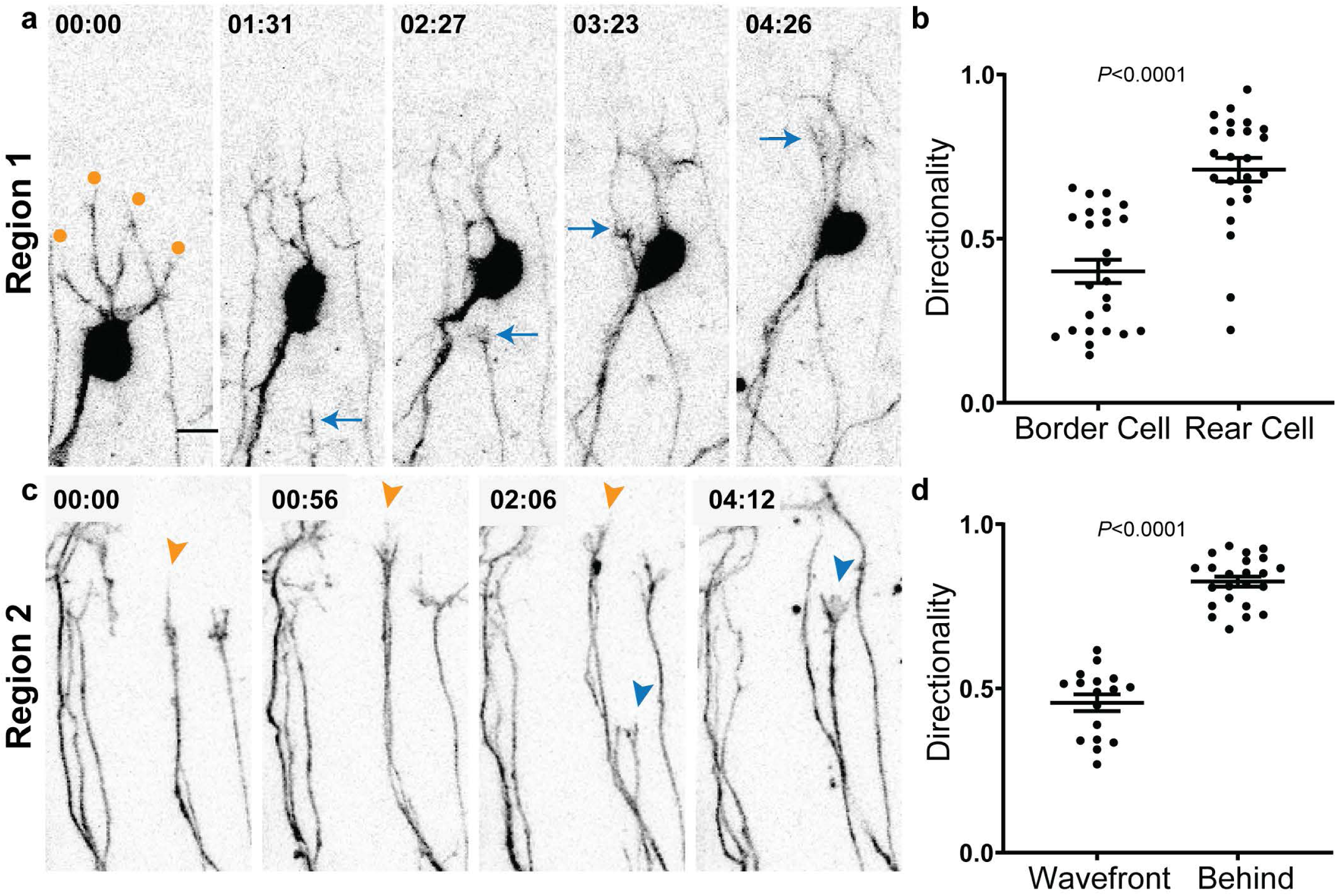
Peripheral process outgrowth behavior differs even with similar regions. (**a-c**) Frames from movies of sparsely labeled tdTomato+ SGNs from *Neurog1^CreERT2^;Ai14* animals as they grow in region 1 (a) (N=3 cochleae) or region 2 (c)(N=3 cochleae). In Region 1 (a), a border SGN extends highly branched processes (orange dots) whereas a peripheral process from a rear cell (cell body out of view) (blue arrows) follows a rapid and directed path. Quantification of 25 border and 24 rear cells confirmed that peripheral processes from rear cells are significantly more directed (b). In Region 2 (c), an SGN process at the wavefront (orange arrowheads) extends many branches and makes little progress. An SGN process arriving from behind (blue arrowheads) moves rapidly and is capped by a large, unbranched growth cone. Quantification revealed significantly more directed growth of processes behind the wavefront (n=17 wavefront processes and 23 trailing processes). Note that not all SGN processes are labeled, so it is not possible to detect fasciculation events reliably. Significance was assessed using an unpaired, two-tailed t-test; *P* values are indicated on the panels. Time is indicated in hh:mm. Scale bar = 10 µm (e). See Videos 5-6.

Analysis of outgrowth behavior in R2 revealed additional heterogeneity here, as well. In this case, the processes had grown too far away from the ganglion to be able to link them definitively to border or rear SGNs. Instead, we compared the behavior of those processes at the wavefront to those that grew up from behind (Fig. 5c and Video 6). We found that in this region, the processes at the wavefront (orange arrowheads, Fig. 5c) were significantly less directed than the processes behind them (blue arrowheads, Fig. 5c) (Fig. 5d) (P<0.0001 by unpaired, two-tailed t-test; n=17 wavefront neurites and 23 follower neurites from N=3 cochleae). At this point in the trajectory, it seems that the trailing processes have fasciculated with the pioneers, which are still navigating forward through the mesenchyme. It was not possible to quantify the degree of fasciculation, as only a subset of processes are labeled. However, it is already established that fasciculation occurs in this region of the mesenchyme ^21^. Thus, within both R1 and R2, SGN processes use different cellular mechanisms to approach the same targets, with some processes seeming to interact with non-neuronal cells in the environment and others relying on fasciculation with other SGN processes that have already forged the way.

### Early SGN processes interact actively with glia that migrate ahead of them

Next, we sought to determine how interactions with glia might contribute to the variation in outgrowth behaviors that we have observed. Several observations suggest that glia may encourage innervation by the earliest processes and hence create a scaffold for later arriving processes. First, the glial cells are present ahead of the wavefront of process outgrowth (Fig. 2d,g). Second, the processes from the rear SGNs are closest to this glial carpet (Fig. 2d’e and Fig. 3b”) and also grow in a faster and more directed manner here (Fig. 5a). Third, when glia are depleted from the cochlea, radial bundle formation is severely disrupted ^25, 26^. Although axon-axon fasciculation is a well-established mechanism for nervous system wiring, the potential influence of neuron-glia interactions is not well understood.

To learn more about how interactions with glia might affect axon growth, we simultaneously imaged SGN peripheral processes and glia in E14-E14.5 cochleae from *Neurog1^CreERT2^;Ai14;PLP-GFP* animals (N=4), characterizing both the interactions at the wavefront and behind, where a scaffold is already in place. We found that the PLP-GFP+ glia migrate in radial chains through the mesenchyme (Fig. 6a). Within the chains, the SGN peripheral processes sometimes grew directly on the glia, such that tdTomato and GFP signals co-localized, but at other times, the tdTomato and GFP signals did not overlap. Quantification revealed that the peripheral processes that contacted glia subsequently grew in a directed manner, often following the glial cell (Fig. 6b). By contrast, SGN processes that were not contacting glia were more exploratory, with a significantly lower direction index (Fig. 6c). Moreover, there were several cases where SGN neurite growth was predicted by a similar change in glial behavior (Videos 7-11). For example, we observed multiple examples where an SGN peripheral process retracted (*, Fig. 6d, g) shortly after the associated PLP-GFP positive glial process had retracted, such that the SGN and glial processes ended up once again on top of each other. Likewise, forward movement of a PLP-GFP process was rapidly followed by forward movement of the associated SGN peripheral process in the same direction (+, Fig. 6e). In other cases, the SGN processes simply grow along pre-existing glial tracts (arrowheads, Fig. 6f,g). This may be especially true behind the wavefront of growth, where the glia seem less active and may instead provide a permissive substrate. Thus, SGN peripheral process outgrowth appears to be facilitated by interactions with glia that are growing in the same direction.

**Figure 6:**
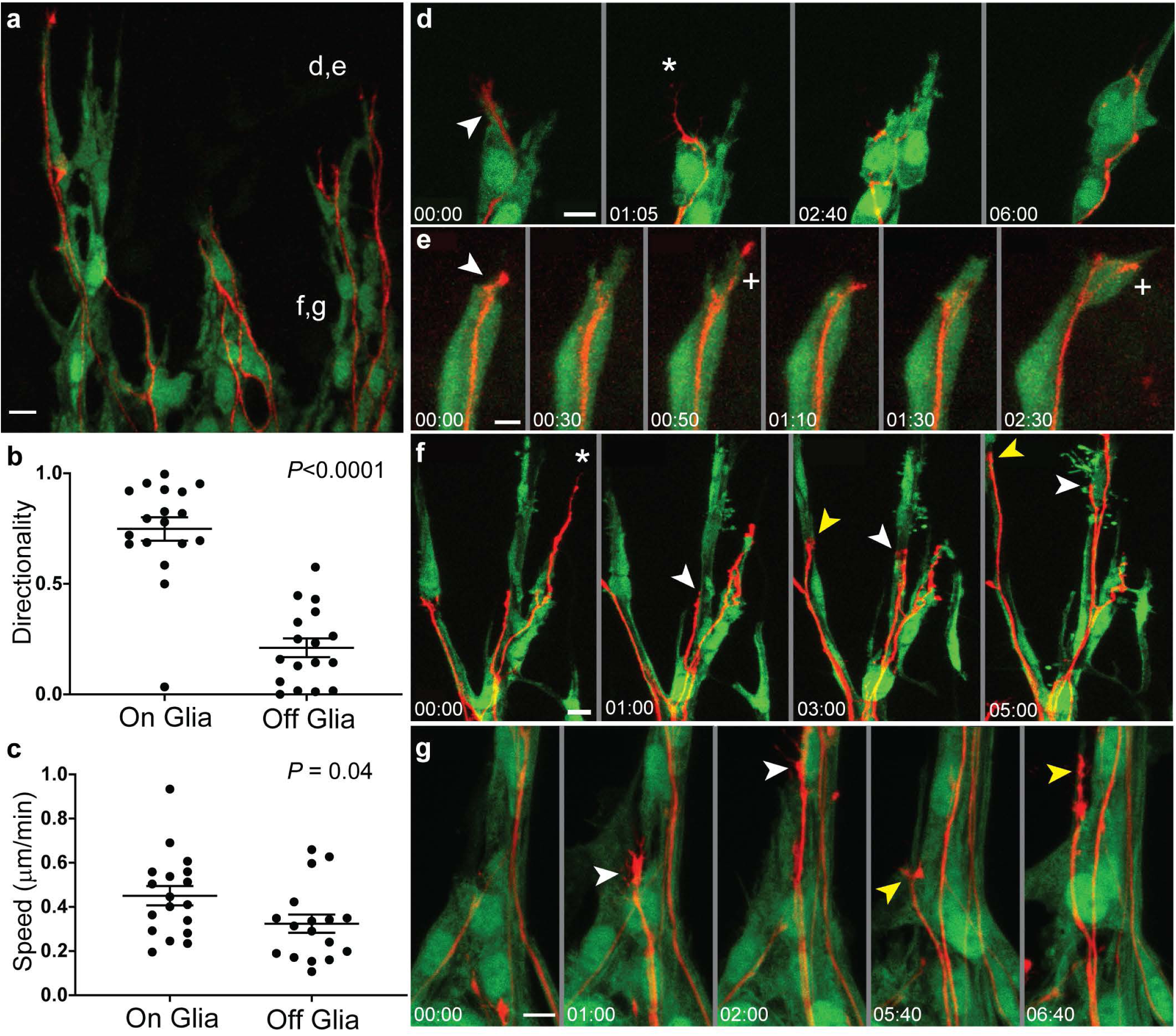
SGN peripheral process behavior is preceded by similar changes in glial outgrowth. (**a**) A low power view of the wavefront of peripheral process outgrowth in a live E14 cochlea from a *Neurog1-^CreERT2^;AI14;PLP-GFP* animal, with individual SGN peripheral processes in red and all glia in green. SGN processes grow through chains of migrating glia that are separated by mesenchyme (not labeled). Time lapse imaging of neuron-glia interactions was performed, focusing either close to the wavefront (as in d and e) or behind (as in f). (**b,c**) Quantification of directionality (b) and speed (c) for neurites that were growing either on or off a glial cell. Processes were more directed when growing on glia. 35 neurites were analyzed from 4 cochleae. Significance was assessed using an unpaired, two-tailed t-test; *P* values are indicated on the panels. (**d,e**) Frames from two movies of SGN outgrowth at the wavefront, with time shown in hh:mm. In one movie (d, Video 7), the distal tip of the SGN process (arrowhead, 00:00) is initially on top of process from a glial cell. By 01:05, the glial cell process has retracted (*). Subsequently, the SGN process also retracts (2:40) and then re-aligns along the second, remaining glial process (06:00). In the second movie (Video 9), an SGN process starts on top of a glial cell (arrowhead, 00:00). Subsequently, the glial cell extends forward (00:30), followed quickly by a branch from the SGN process (+, 00:50). After a quick retraction (01:10), the glial process again extends forward (01:30), followed by the SGN neurite (+, 2:30). (**f**) A lower power movie of neuron-glia interactions behind the wavefront (Video 10) illustrates multiple examples of SGN neurite behavior, including a retraction (*) and two examples of fasciculation along glial tracts (white and yellow arrowheads). (**g**) Frames from a movie of SGN outgrowth behind the wavefront (Video 11). Within a chain of glia, one SGN peripheral process capped by a large growth cone moves straight forward in a short period of time (white arrowhead, 01:00 to 02:00). Subsequently, a second process (yellow arrowhead, 05:40 to 06:40) makes similarly rapid and directed process, possibly fasciculating along a thin PLP-GFP+ process ahead of it. See also Video 8 for an additional example of neuron-glia interactions. Scale bars = 10 µm.

## Discussion

Although there is abundant evidence that axons can be directed to their targets by a combination of positive and negative cues, the full range of behaviors that ensure reliable axon guidance *in vivo* remains to be defined. We examined this question by characterizing outgrowth in the cochlea, where axons grow through a heterogeneous environment to reach their targets in the sensory epithelium. In this unique cellular landscape, processes exhibit different patterns of outgrowth depending on when they are born as well as their position within the ganglion. For instance, during the initial stages of cochlear wiring, SGNs in the rear of the ganglion extend processes that have easy access to a glial carpet *en route* to the organ of Corti. The processes grow faster and make fewer changes in direction in this region. Follower SGNs closer to the border of the ganglion, on the other hand, must grow down towards this path, where they are able to fasciculate with the processes from the rear. Moreover, imaging studies suggest that the glia may play an active role in eliciting and directing axon outgrowth towards the organ of Corti, as well as providing an attractive, permissive surface. Together, our findings support a model in which glia pattern an axon scaffold and therefore enable rapid and directed growth of SGN peripheral processes towards the organ of Corti. Such a mechanism provides a simple way to establish the basic topography of the cochlea. Rather than sensing cues that tell them where they are along the apical-basal axis, individual SGN peripheral processes may simply need to follow a straight path towards the organ of Corti, with growth encouraged by glia and further corralled by surrounding mesenchyme ^21^. Additionally, with the flexibility to grow either along glia or along each other, SGN neurites can navigate towards their target reliably regardless of where they are positioned within the three dimensional structure of the ganglion.

Although it is now well appreciated that astrocytes and microglia directly impact synapse formation ^33^, our findings highlight an even earlier role for glia during axon outgrowth and guidance. We find that in the cochlea, glia are not only along the migratory path, but are in fact actively extending ahead of the SGN processes, forming chains that presage the appearance of radial bundles a day later. Moreover, individual SGN processes change their behavior upon contacting the glia, such that processes that are on glia grow faster and make fewer changes in direction compared to those that are off the glia. Thus, the SGN processes that are closest to the glial carpet, simply due to their position in the ganglion, can serve as pioneers for those that arrive later and thus encounter a different cellular environment.

Our observations support the classic idea that glial-guided growth of pioneer axons may be a general mechanism for establishing scaffolds throughout the nervous system. In fact, forty years ago, Singer et al. put forward the idea of a glial “blueprint” of the nervous system based on the close proximity of glial precursors and growth cones in the Newt spinal cord ^34^. Likewise, in the cortex, glial processes form a “sling” straddling the lateral ventricles ^35^. Similar to what we observed by imaging processes as they grow in the cochlea, glial processes appear to precede pioneer axons in the cortex as they cross the midline, with follower axons fasciculating. Additional evidence for glial-guided growth has come from genetic ablation of glia in flies, followed by analysis of longitudinal axons ^36^. As in the cochlea, pioneer axons extend towards glia. When the glia are absent, the growth cones move more slowly and some axons show altered fasciculation. A role in fasciculation has also been shown in studies of the developing lateral line in zebrafish ^37^, although in this case the glia seem to migrate along the axons.

Although the coincidence of glia and early axons is suggestive, it has been difficult to figure out exactly how the glia contribute to axon growth. When the glial sling is lesioned, the corpus callosum fails to form ^35^. However, this does not mean that the glia instructed formation of the corpus callosum, as it is equally plausible that they play a permissive role. One challenge is the fact that glia also provide trophic support for neurons, so complete depletion of the glia can cause massive changes in neuron number and organization, making it hard to know whether any early axon guidance phenotypes are primary or secondary ^38^. Local ablation of glia, for instance in developing chicken embryos, offers one potential solution and has revealed minor axon guidance defects ^24^. However, in these experiments, surrounding glia can proliferate and rescue the ablated area, again complicating interpretations. More definitive evidence that glia guide early axons has come from C. elegans, where glia are not required for neuronal survival ^39^. Here, specific molecules have been identified, further emphasizing the need for molecular instructions from the glia that are read out by the extending pioneer axons. An important next step will be to find the molecules that might mediate neuron-glia interactions in the cochlea.

The use of glia to organize early axon outgrowth may complement canonical guidance systems and thereby improve the fidelity of circuit assembly. There is ample evidence that isolated axons can respond to instructive cues, so a role for glia is unlikely to be required for all guidance decisions. Instead, glia may collaborate with these guidance systems, perhaps both permissively and instructively. One possibility is that the glia are responding to the same cues and simply provide a substrate that is more permissive for axon outgrowth. In this model, axon guidance errors are reduced by taking advantage of tiered levels of fasciculation: the early glia pave the initial path for the pioneer axons, which in turn provide a surface for follower axons. This way, the later axons are more likely to find the right path, even as both the distances and cellular complexity increase. Testing such a model is not straightforward as one would not expect axons to become completely lost but instead to be slightly slower or make more mistakes *en route*. Identifying these types of subtle guidance defects will require analysis of individual axon trajectories and more careful quantification, similar to how a role for floor plate-derived Netrin-1 was demonstrated ^8, 9^. It is also possible that the glia themselves provide additional synergizing cues that directly impact neurite outgrowth behavior. Glia do in fact express classic guidance cues, such as Semaphorins ^40^, and it will be interesting to test whether glial-derived guidance cues play any role during cochlear wiring in the future. Conversely, it will be important to learn how glia themselves are guided towards their targets.

Our findings have interesting implications for efforts to re-wire the nervous system after damage. It has long been known that peripheral glia permit axon regeneration, whereas central glia are inhibitory ^41^. Our work suggests that peripheral glia are not only be permissive, but might in fact be able to actively encourage axon re-extension. In the cochlea, SGN processes lose their synapses and retract following exposure to excessively loud sounds in animal models ^42^. Similar loss of synapses has been observed in aged human cochlear specimens, suggesting that damage to SGN processes accumulates over a life time of noise exposure and hence contributes to age-related hearing loss ^43, 44^. However, SGN peripheral processes persist despite being disconnected from the hair cells. One exciting possibility is that the Schwann cells that remain could be used to stimulate re-extension to the hair cells. Importantly, Schwann cells are known to be unusually plastic and can de-differentiate to produce new cells in other regions of the nervous system ^45^. Thus, it is possible that the cochlea could be re-wired by finding ways to de-differentiate the Schwann cells so that they regain their youthful outgrowth abilities.

## Materials and Methods

### Animals

Bulk labeling of SGNs was accomplished by crossing *Bhlhb5^Cre^* mice ^46^ to a Cre-dependent tdTomato reporter strain (*Ai14*; #007908 Jackson Laboratory) ^47^. SGNs were sparsely labeled by crossing *Neurog1^CreERT2^* mice ^30^ to the same reporter strain. Mice harboring a GFP reporter for PLP promoter activity (“*PLP-GFP*”) were used to label cochlear glia ^32^. Animals were PCR genotyped using primers for Cre, tdTomato, or GFP; embryos were genotyped using a fluorescent dissecting microscope. To obtain timed pregnancies, male mice heterozygous for *PLP-GFP*, *Ai14*, and an appropriate *Cre* strain were bred with adult CD1 females (Charles River Laboratories). Noon on the day of the plug was considered embryonic day 0 (E0). Embryos and pups of either sex were used. Animals were maintained and handled according to protocols approved by the I.A.C.U.C. at Harvard Medical School.

### Organ Culture and Live Imaging

Cochlear explants were collected and cultured as described ^19, 48^. In brief, fetuses were isolated from timed pregnant females, checked for fluorescence, and then dissected. Cochleae were placed into an imaging chamber, centered on a 1 mm hole punched into a small square of electrostatically charged cellulose membrane and held in place with a vitaline tissue drape. The imaging chamber contained culture media (defined DF12, Glutamax (Gibco), N2 supplement, 25mM Glucose, 20mM Hepes, 25mM sodium bicarbonate, and 5% FBS (Gibco), no antibiotics are used) that was closed to atmosphere and heated to maintain the reported internal temperature at 37 °C. Cochleae were imaged along the mid-base region for each experiment. Using an Olympus FluoView 1000 confocal microscope, images were acquired every 7-14 minutes with a 60x oil lens through the entire area focal depth of interest (50-100 µm) in z-steps of 2-5 µm. Time-lapse files were tracked and analyzed with MetaMorph (Molecular Devices) and ImageJ software (NIH).

Each time-lapse file was converted into RGB-depth coded movies to visualize individual neurite projections along the z-axis. SGN processes were manually tracked and defined as any and all protrusions peripheral from the ganglion that could be delineated from its neighbors and could be tracked for more than four time intervals. DiPER code ^49^ was used to analyze track speed and directionality. The XY coordinates for all tracks were recorded and plotted to show their orientation and start/stop points using Prism (Graphpad).

Regions of interest were defined by measuring the position of the neurites relative to the border of the ganglion, delineated by the SGN cell bodies positioned there. Neurites in R1 were from ∼E14.25-.5 cochleae and < 50 µm from the border. R2 neurites were imaged in ∼E15 cochleae and were 50-100 µm from the ganglion. R3 neurites were imaged in ∼E15.5 cochleae and were >100 µm from the ganglion. Within the R1 and R2 regions, “wavefront” neurites were defined as those most distally extended at the time of imaging. Any neurites >10 µm modiolar to these distalmost neurites were classified as “behind.”

The behavior of SGN neurites on and off glia was quantified by tracking neurites as described above while simultaneously monitoring the position of PLP-GFP labeled glia in a separate channel. “On” SGN neurites occupied a position that could not be resolved from the PLP-GFP signal in the glia. “Off” SGN neurites, by contrast, extended >1 µm away from PLP-GFP+ glia for >2 time points.

### Immunostaining

The following primary antibodies and dilutions were used in this study: rabbit anti-DsRed (Clontech; 1:1000), Goat anti-Sox10 (Santa Cruz; 1:500), Rabbit anti-Tuj (Biolegend, 1:1000), Goat anti-GFP FITC (Abcam 1:500), Chicken anti-Neurofilament (Millipore, 1:1000) and Chicken anti-GFP (Aves, 1:2000). Alexa-conjugated secondary antibodies raised in donkey were all used at the same concentration (Jackson ImmunoResearch; 1:500). For immunostaining, tissues were dissected, washed twice with PBS, and fixed with 4% paraformaldehyde (PFA) in PBS overnight at 4°C, and then washed three times in PBS.

For wholemount immunostaining, dissected cochleae were permeabilized and blocked at room temperature for 2 hrs in phosphate buffer saline (PBS) containing 0.3% Triton, 3% Bovine Serum Albumen, and 0.01% sodium azide. Primary antibody hybridization was performed in the same blocking buffer solution overnight at 4°C, followed by three washes in PBS at room temperature. Secondary antibody hybridization was performed in blocking buffer overnight at 4°C and extensively washed in PBS at room temperature the following day. Prior to imaging, tissues were cleared in BABB (Benzyl alcohol:Benzyl Benzoate, 1:2 ratio). Briefly, cochlea were transferred through a graded series into 100% methanol and then placed flush on a glass slide within a rectangular silicon grease reservoir. Methanol within the reservoir was gently aspirated and replaced with a 1:1 solution of methanol:BABB. After partial clearing, this 1:1 solution was replaced with 100% BABB and the silicon grease reservoir was sealed with a glass coverslip.

For section immunostaining, fixed E14-E14.5 heads were cryoprotected by incubating in 10% and 20% sucrose in PBS for one night each and then in 30% sucrose in NEG-50 (Richard-Allan Scientific) overnight, all at 4° C. Heads were embedded in NEG-50 and stored at −80° C prior to sectioning at 20 µm thickness. After sectioning, slides were allowed to dry at −80° C overnight. Antigen retrieval in 10 mM citrate buffer (pH 6.0) was done for 20 min before commencing with the staining protocol. Blocking and antibody hybridization steps were carried out in PBS containing 5% Normal Donkey Serum and 0.5% Triton. Slides were blocked at room temperature for one hour, followed by primary antibody hybridization overnight at 4° C. Hybridization of the secondary antibody was carried out for one hour at room temperature before washing thoroughly with PBS, incubating in DAPI and mounting a coverslip.

### Fixed tissue analysis

After immunostaining and clearing, wholemount *Neurog1^CreERT2^;Ai14* cochleae (N=7) were imaged on an Olympus FluoView 1000 confocal microscope. The entire volume of the mid-basal region of the ganglion was imaged in consecutive z-slices separated by 0.5 µm. These volumes were then rendered using Imaris. Neurons that could be clearly differentiated from adjacent cells and that could be viewed all the way from the cell body to the distalmost neurites were reconstructed using a combination of manual and automated methods that defined the outline of the cell plane by plane. The defined regions of each plane were then combined to create three-dimensional rendered surfaces. For analysis, the surfaces representing the cells of each region were subdivided by measuring the distance from the border to the rear of the cell group and dividing into three equal parts. Border (n=28), mid (n=52), and rear cells (n=71) were assigned to groups based on the position of the center of the cell body. The primary process of each cell was considered to be the one which extended the greatest length from the cell body towards the organ of Corti. Imaris software was used to analyze each rendered cell to determine neurite length, defined as the total distanced covered; distance, defined as the linear distance between the neurites’s origin and endpoint; and directionality, defined as distance/length; z-displacement, defined as the distance a process travelled along the z axis between stacks; two slope values, calculated as z-displacement either over length or over distance; branch points, defined as the number of places where a branch of at least 5 μm splits off; and distance to branch, which is defined as the length from the process’s endpoint to the nearest branch of at least 5 μm. Measurements were taken using a series of fixed points along the surfaces. The average and SEM for each parameter was found for each of the three categories so that the characteristics of cell processes could be compared between regions. Only measurements for length, directionality, and slope are presented. Data from each of the seven cochleae are shown independently in Supplemental Figure 2.

For analysis of neuron-glia interactions, wholemount *Neurog1^CreERT2^;Ai14;PLP-GFP* cochleae that had been stained for GFP, tdTomato and/or Tuj were cleared and then imaged on an Olympus FluoView 1000 confocal microscope, followed by analysis using Imaris. TdTomato+ or TuJ1+ neurites in R1 were identified and then assessed for proximity to a PLP-GFP+ glial cell, simply noting whether the neurite was ahead of the glial cell or behind it.

### Statistics

Sample sizes were determined without any expectation of the effect size, but with cochleae from at least three different animals. All cells that could be confidently reconstructed or scored were included. One cochlea was excluded from the analysis in Figure 3 due to incomplete labeling that prevented an unbiased assessment of morphology. SGN reconstructions were analyzed by an independent investigator blind to the possible outcomes. All statistical analyses were done using Prism software (Graphpad). Data shown in Figures 2 and 3 were first assessed for normality and then analyzed using ANOVA with Tukey’s multiple comparison test. Data in Figure 4 were analyzed with multiple t-tests using the Holm-Sidak method. Data in Figures 5-7 were analyzed using an unpaired two-tailed t-test. Results with a *P*<0.05 were considered statistically significant. Sample sizes and *P* values are reported in the Figure Legends and are summarized in Supplemental Table 1.

## Acknowledgements

We thank D. Corey (Department of Neurobiology, Harvard Medical School) for providing access to his confocal microscope. We are also grateful to members of the Goodrich laboratory and Dr. Maxwell Heiman for helpful conversations and comments on the manuscript.

No competing interests declared.

---

**Video 1: Sparse labeling of SGNs reveal heterogeneous morphology within the E14 ganglion**. A video illustrating a typical set of reconstructed SGNs in the ∼E14 *Neurog1^CreERT2^; Ai14* cochlea. Rear SGNs are shown in purple; middle SGNs in green; and border SGNs in yellow. The same SGNs are shown in Fig. 3b.

**Video 2: SGN peripheral process outgrowth in R1.** An example of SGN peripheral process behavior at the wavefront of early outgrowth, as imaged in a ∼E14.25 *Bhlhb5^Cre^; Ai14* cochlea. This region corresponds to R1, as illustrated in Fig. 4a, b. Frames from this movie are shown in Fig. 4g.

**Video 3: SGN peripheral process outgrowth in R2.** A representative example of SGN peripheral process behavior in an E15 *Bhlhb5^Cre^; Ai14* cochlea. By this time, the wavefront of outgrowth has reached R2, as illustrated in Fig. 4 a, b.

**Video 4: SGN peripheral process outgrowth in R3.** A representative example of SGN peripheral process behavior in an E15.5 *Bhlhb5^Cre^; Ai14* cochlea. By E15.5, peripheral processes have reached R3, where the organ of Corti is developing, as illustrated in Fig. 4a, b.

**Video 5: Individual SGN processes follow distinct trajectories in R1.** A movie of SGN peripheral process behavior in R1 of an E14.25 *Neurog1^CreERT2^; Ai14* cochlea. In this case, a random subset of SGNs are fluorescently labeled, making it possible to distinguish processes from SGNs whose cell bodies sit at the border from those whose cell bodies are further behind and out of view (movement of the growth cone is indicated by arrows). Frames from this movie are shown in Fig. 5a.

**Video 6: Individual SGN processes exhibit distinct outgrowth behaviors within R2**. A movie of SGN peripheral process behavior in R2 of an ∼E15 *Neurog1^CreERT2^; Ai14* cochlea. By this stage, the processes at the wavefront of growth show more exploratory behavior than those that start from further behind. Frames from this movie are shown in Fig. 5b.

**Video 7-11: Neuron-glia interactions in the developing cochlea.** 5 movies from ∼E14-E14.5 *Neurog1^CreERT2^; Ai14; PLP-GFP* cochlea, with tdTomato+ SGN peripheral processes in red and PLP-GFP+ glia in green. Videos 7-9 highlight behavior at the wavefront; Videos 10 and 11 illustrate behavior behind the wavefront. Frames from Video 7 are shown in Fig. 6d; from Video 9 in Fig. 6e; from Video 10 in Fig. 6f; and from Video 11 in Fig. 6g. Video 8 is not shown elsewhere but is included as an additional example of neuron-glia interactions.

**Supplemental Figure 1:**
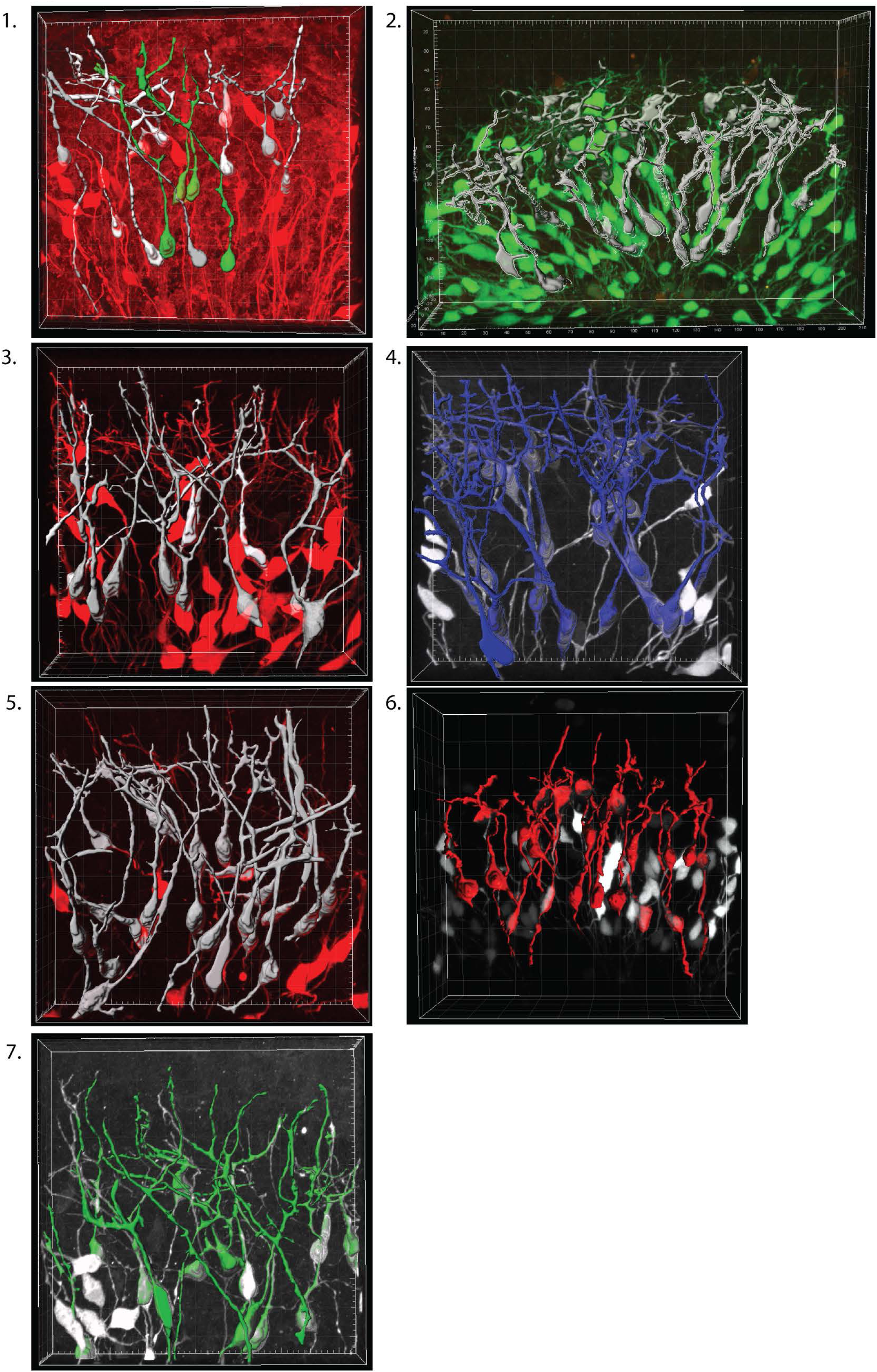
Snapshots of the seven reconstructed cochleae used to generate the data shown in Figure 3.

**Supplemental Figure 2:**
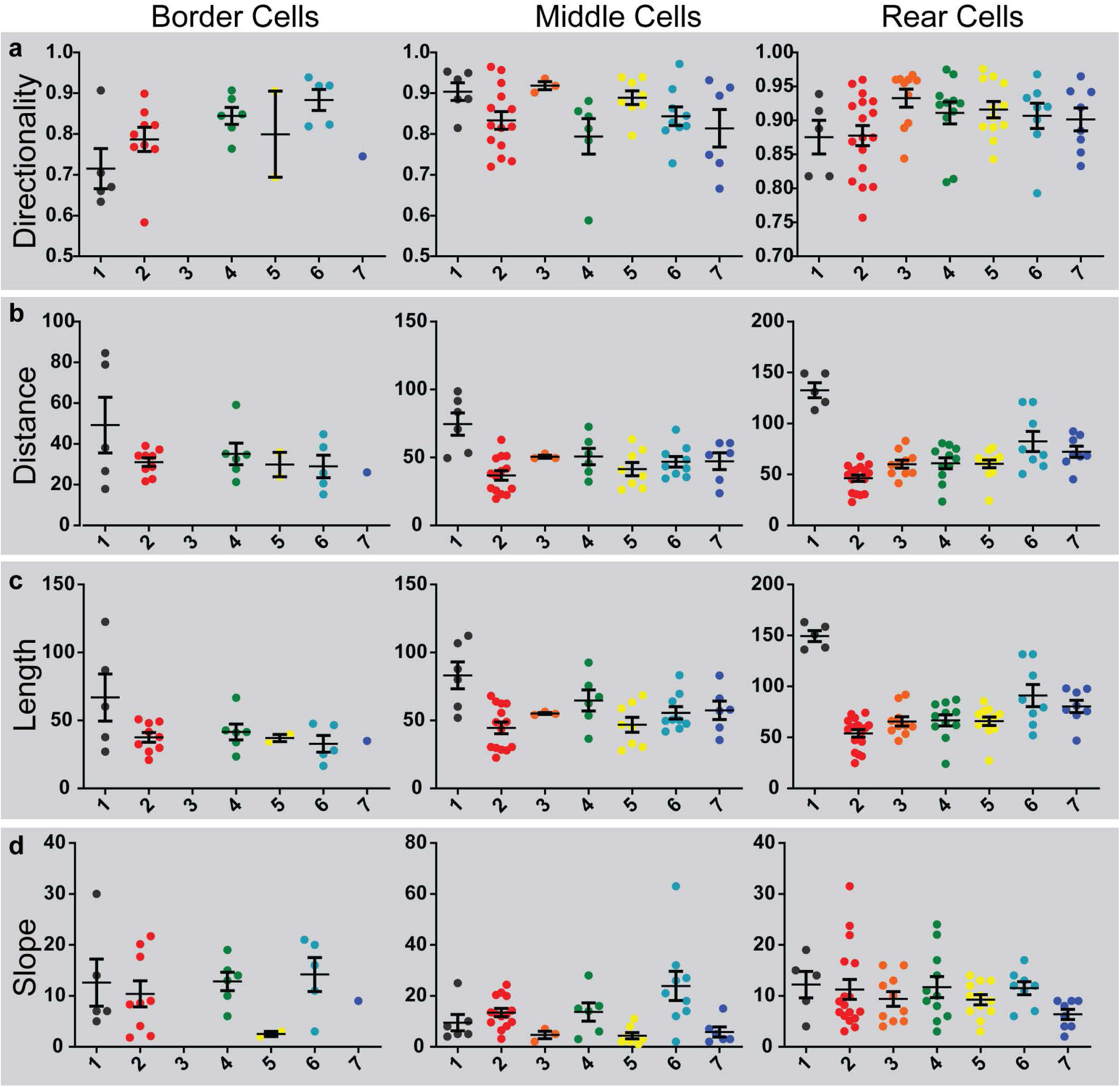
Metrics from the seven individual cochlea used to generate data in Figure 3, numbered as shown in Supplemental Figure 1. For each cochlea, directionality (a), distance (b), length (c), and slope (d) were calculated for cells sitting in the border, middle or rear regions. Please note that the axes change for each population so that all of the individual data points can be seen easily. Mean and standard error of mean are shown for each population.

**Supplemental Table 1:**
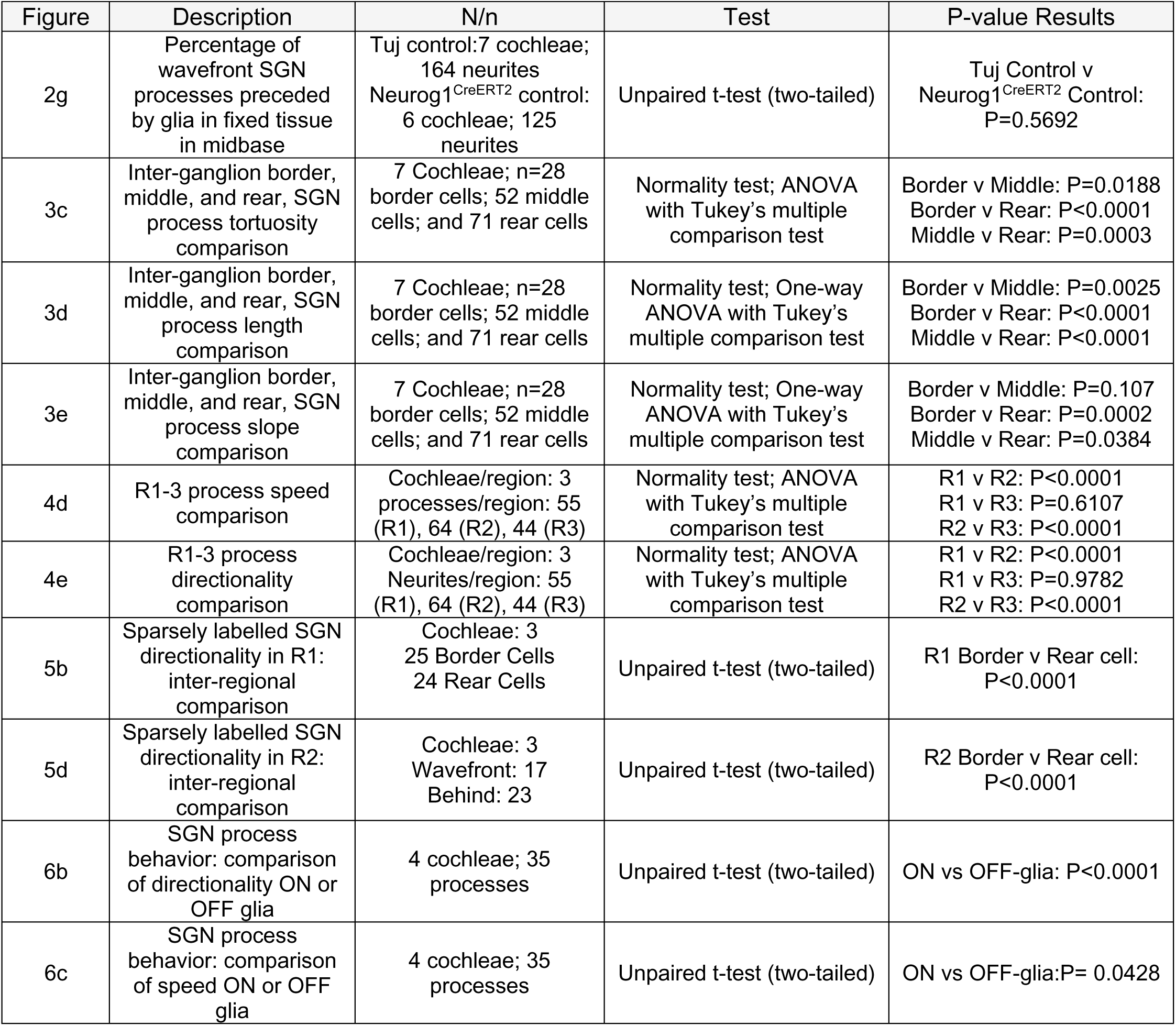
A summary of the group sizes and statistical tests used to analyze data presented in this manuscript.

| Figure | Description | N/n | Test | P-value Results |
| --- | --- | --- | --- | --- |
| 2g | Percentage of wavefront SGN processes preceded by glia in fixed tissue in midbase | Tuj control: 7 cochleae; 164 neurites<br>Neurog1<sup>CreERT2</sup> control: 6 cochleae; 125 neurites | Unpaired t-test (two-tailed) | Tuj Control v Neurog1<sup>CreERT2</sup> Control: P=0.5692 |
| 3c | Inter-ganglion border, middle, and rear, SGN process tortuosity comparison | 7 Cochleae; n=28 border cells; 52 middle cells; and 71 rear cells | Normality test; ANOVA with Tukey's multiple comparison test | Border v Middle: P=0.0188<br>Border v Rear: P<0.0001<br>Middle v Rear: P=0.0003 |
| 3d | Inter-ganglion border, middle, and rear, SGN process length comparison | 7 Cochleae; n=28 border cells; 52 middle cells; and 71 rear cells | Normality test; One-way ANOVA with Tukey's multiple comparison test | Border v Middle: P=0.0025<br>Border v Rear: P<0.0001<br>Middle v Rear: P<0.0001 |
| 3e | Inter-ganglion border, middle, and rear, SGN process slope comparison | 7 Cochleae; n=28 border cells; 52 middle cells; and 71 rear cells | Normality test; One-way ANOVA with Tukey's multiple comparison test | Border v Middle: P=0.107<br>Border v Rear: P=0.0002<br>Middle v Rear: P=0.0384 |
| 4d | R1-3 process speed comparison | Cochleae/region: 3 processes/region: 55 (R1), 64 (R2), 44 (R3) | Normality test; ANOVA with Tukey's multiple comparison test | R1 v R2: P<0.0001<br>R1 v R3: P=0.6107<br>R2 v R3: P<0.0001 |
| 4e | R1-3 process directionality comparison | Cochleae/region: 3 Neurites/region: 55 (R1), 64 (R2), 44 (R3) | Normality test; ANOVA with Tukey's multiple comparison test | R1 v R2: P<0.0001<br>R1 v R3: P=0.9782<br>R2 v R3: P<0.0001 |
| 5b | Sparsely labelled SGN directionality in R1: inter-regional comparison | Cochleae: 3<br>25 Border Cells<br>24 Rear Cells | Unpaired t-test (two-tailed) | R1 Border v Rear cell: P<0.0001 |
| 5d | Sparsely labelled SGN directionality in R2: inter-regional comparison | Cochleae: 3<br>Wavefront: 17<br>Behind: 23 | Unpaired t-test (two-tailed) | R2 Border v Rear cell: P<0.0001 |
| 6b | SGN process behavior: comparison of directionality ON or OFF glia | 4 cochleae; 35 processes | Unpaired t-test (two-tailed) | ON vs OFF-glia: P<0.0001 |
| 6c | SGN process behavior: comparison of speed ON or OFF glia | 4 cochleae; 35 processes | Unpaired t-test (two-tailed) | ON vs OFF-glia: P= 0.0428 |

